# A lineage-tracing tool to map the fate of hypoxic tumour cells

**DOI:** 10.1101/2020.03.03.974857

**Authors:** Jenny A.F. Vermeer, Jonathan Ient, Bostjan Markelc, Jakob Kaeppler, Lydie M.O. Barbeau, Arjan J. Groot, Ruth J. Muschel, Marc A. Vooijs

**Author notes:** Authors contributed equally (Jenny A.F. Vermeer, Jonathan Ient).

## Abstract

Intratumoural hypoxia is a common characteristic of malignant treatment-resistant cancers. However, hypoxia-modification strategies for the clinic remain elusive. To date little is known on the behaviour of individual hypoxic tumour cells in their microenvironment. To explore this issue in a spatial and temporally-controlled manner we developed a genetically encoded sensor by fusing the O_2_-labile Hypoxia-Inducible Factor 1α to eGFP and a tamoxifen-regulated Cre recombinase. Under normoxic conditions HIF-1α is degraded but under hypoxia, the HIF-1α-GFP-Cre-ER^T2^ fusion protein is stabilised and in the presence of tamoxifen activates a tdTomato reporter gene that is constitutively expressed in hypoxic progeny. We visualise the random distribution of hypoxic tumour cells from hypoxic or necrotic regions and vascularised areas using immunofluorescence and intravital microscopy. Once tdTomato expression is induced, it is stable for at least 4 weeks. Using this system, we could show that the post-hypoxic cells were more proliferative *in vivo* than non-labelled cells. Our results demonstrate that single-cell lineage tracing of hypoxic tumour cells can allow visualisation of their behaviour in living tumours using intravital microscopy. This tool should prove valuable for the study of dissemination and treatment response of post-hypoxic tumour cells *in vivo* at single-cell resolution.

**Summary Statement:** Here we developed and characterised a novel HIF-1α-Cre fusion gene to trace the progeny of hypoxic tumour cells in a temporal and spatially resolved manner using intravital microscopy.

## Introduction

Many solid tumours contain areas of hypoxia, which is the result of O_2_ demand (rapid proliferation) exceeding O_2_ supply (aberrant vasculature) (Thomlinson and Gray, 1955; Brahimi-Horn, Chiche and Pouysségur, 2007). Due to limits on O_2_ diffusion from the blood vessels, tumours experience chronic hypoxia and necrosis in regions distant from the vasculature. Acute or cycling hypoxia also occurs in tumours due to temporary occlusion of blood vessels obstructing perfusion leading to areas of hypoxia, which become re-oxygenated when the obstruction is relieved (Dewhirst, 2009; Salem *et al.*, 2018). Tumour hypoxia is strongly associated with a worse outcome in many different cancers irrespective of treatment (Vaupel and Mayer, 2007) and tumour hypoxia is a direct factor in resistance to some chemotherapy and radiation therapy. Direct O_2_ measurements in clinical studies using oxygen needle electrodes indicate hypoxia is strongly associated with local regional control in head and neck squamous cell carcinoma (Nordsmark *et al.*, 2005), prostate (Milosevic *et al.*, 2012) and cervix (Fyles *et al.*, 1998) cancer patients treated with radiotherapy. Hypoxia imaging using PET tracers such as ^18^F-Hx4, ^18^F-MISO, and ^18^F-FAZA have demonstrated strong associations of tracer uptake with outcome (Lehtiö *et al.*, 2004; Dubois *et al.*, 2011; Fleming *et al.*, 2015).

The main adaptive response to hypoxia is the stabilisation of the oxygen regulated hypoxia inducible factor alpha (HIF-α) proteins HIF-1α HIF-2α and HIF-3α. HIF-α proteins are predominantly regulated post-translationally through oxygen dependent prolyl and asparagine hydroxylases, which hydroxylate specific proline and asparagine residues on the HIF-α oxygen dependent degradation (ODD) domain. Hydroxylation of asparagine inhibits the recruitment of the transcriptional regulator p300. Prolyl hydroxylation promotes interaction with the von Hippel-Lindau (pVHL) protein, which recruits an E3 ubiquitin ligase, targeting HIF-α for proteasomal degradation (Ivan *et al.*, 2001). Under hypoxic conditions, the activity of prolyl and asparagine hydroxylases is attenuated leading to accumulation of HIF-α proteins. HIF-α then translocates to the nucleus where it binds the constitutively expressed HIF-1ß protein and the co-activator p300. The HIF transcriptional complex is known to trans-activate over 1,500 target genes through binding to hypoxia response elements (HREs) located in the target genes or flanking sequences (Prabhakar and Semenza, 2015). HIF target genes play a role in a broad range of pathways including those involved in angiogenesis (VEGF), metabolism (GLUT-1), cell proliferation (TGF-α), cell adhesion (MIC2), pH regulation (CAIX) and cell survival (TGF-α) among others (Jubb, Buffa and Harris, 2010; Wilson and Hay, 2011; Muz *et al.*, 2015; LaGory and Giaccia, 2016). It has long been known that the concentration of oxygen within the tumour correlates with the efficacy of radiotherapy (Gray *et al.*, 1953). HIF-α can be stabilised in relatively mild hypoxia however much more severe hypoxia plays a larger role in radiotherapy resistance due to the decreased effectiveness of radiotherapy at these very low oxygen conditions. Even so, hypoxia and higher levels of HIF are associated with, and their presence also correlates with more aggressive tumours, therapy resistance, immunosuppression, metastasis, and poor prognosis (LaGory and Giaccia, 2016). Elevated HIF-1α and HIF-2α levels have also been shown to be associated with poor prognosis in a number of cancers (Giatromanolaki *et al.*, 2001; Ioannou *et al.*, 2009; Zheng *et al.*, 2013; Ren *et al.*, 2016; Roig *et al.*, 2018).

Because of its strong correlation with adverse patient outcome, hypoxic modification in tumours has been an area of intense basic and translational research and drug development. A systematic review of 10,108 patients across 86 trials that were designed to modify tumour hypoxia in patients that received primary radiotherapy alone showed that overall modification of tumour hypoxia significantly improved the effect of radiotherapy but had no effect on metastasis (Overgaard, 2007).

Accelerated radiotherapy with carbogen and nicotinamide (ARCON) which increases tumour oxygenation to improve radiotherapy treatment has shown limited success in a Phase III clinical trial. ARCON with improved 5 year regional control specifically in patients with hypoxic tumours, however, no improvement in disease free or overall survival was found (Janssens *et al.*, 2012).

The hypoxia activated pro-drug Evofosfamide was studied in a Phase III clinical trial. Evofosfamide improved progression free survival as well as higher objective response rate, however the trial failed as the primary endpoint (overall survival time) was not significantly improved (Van Cutsem *et al.*, 2016). Unfortunately, while a few clinical trials have been successful, many hypoxia modification or targeting trials have failed because of underpowered studies and the lack of hypoxia biomarkers to stratify responders among others (Spiegelberg *et al.*, 2019).

Although many of the molecular mechanisms of how cells respond to hypoxia are known, how the hypoxic cells may contribute to poor prognosis is still poorly understood. It is known that hypoxic cells are more resistant to treatment and more likely to disseminate and develop into metastases (Harada *et al.*, 2012; Muz *et al.*, 2015; Godet *et al.*, 2019). However, a direct demonstration of the cell autonomous phenotypes of hypoxic tumour cells within the primary tumour and their interplay with the tumour microenvironment remains understudied.

Using the ODD domain of the HIF-1α protein as an oxygen sensor fused to a tamoxifen inducible CreER^T2^ recombinase, Harada and colleagues elegantly used lineage tracing of hypoxic cells and their progeny in colon carcinoma xenografts (Harada *et al.*, 2012). They showed that hypoxic cells were able to survive irradiation and formed a large proportion of the recurrent tumour after 25 Gy irradiation. High HIF-1 activity was also found in cells that experience radiation-induced reoxygenation. HIF-1 positive cells after irradiation induced re-oxygenation also translocated towards blood vessels and this translocation was suppressed by HIF inhibitors. Godet et al. recently showed through an alternative hypoxia lineage tracing system, that post-hypoxic tumour cells in mice maintain a ROS-resistant phenotype. This provides a survival advantage in the blood stream therefore promoting their ability to form distant metastases (Godet *et al.*, 2019). One limitation of the aforementioned systems is the relatively long time in continuous hypoxia needed before labelling of cells was achieved, limiting the systems predominantly to areas of sustained chronic hypoxia. These studies also did not visualise individual hypoxic tumour cells within the tumour microenvironment.

In this present study we developed an alternative approach to lineage trace the fate of hypoxic tumour cells that directly reports HIF-1α stabilization rather than the hypoxia transcriptional response. The continuous expression of the system we created allows identification of cells experiencing acute as well as chronic hypoxia and is achieved through a genetically encoded hypoxia sensor composed of a GFP-tagged HIF-ODD-GFP-CreER^T2^ fusion protein herein known as MARCer. Once HIF-1α is stabilised the addition of tamoxifen leads to the cre-mediated activation of a ubiquitously expressed tdTomato, labelling hypoxic cells and their progeny. These fluorescent markers enabled intravital imaging using window chambers (Kedrin *et al.*, 2008), tracing the fate of hypoxic tumour cells at the single cell level within the primary tumour.

## Results

### Hypoxia induces eGFP and tdTomato expression in HIF-MARCer reporter cells

To establish a HIF-cell-tracing method (HIF-MARCer) amenable for in vivo hypoxia imaging, H1299 non-small cell lung carcinoma cells were transduced with hypoxia-inducible factor (HIF)-1α-eGFP-CreER^T2^ cDNA (MARCer fusion protein) expression vector and with a loxP-flanked STOP tdTomato cassette (H1299-MR cells, Fig. 1A). Thus, under hypoxia, the tamoxifen regulated HIF-CRE fusion protein will excise the STOP codons leading to tdTomato expression which will persist under normoxia. To test this system, H1299-MARCer cells were exposed to hypoxia (0.2% O_2_) or Deferoxamine mesylate (DFO, a hypoxia mimetic) *in vitro* resulting in induction of eGFP and HIF-1α protein expression, which was degraded within minutes after re-exposure to normoxia and corresponded with the levels of the endogenous HIF-1α protein (Fig. 1B,C, Fig. S1A).

**Figure 1.**
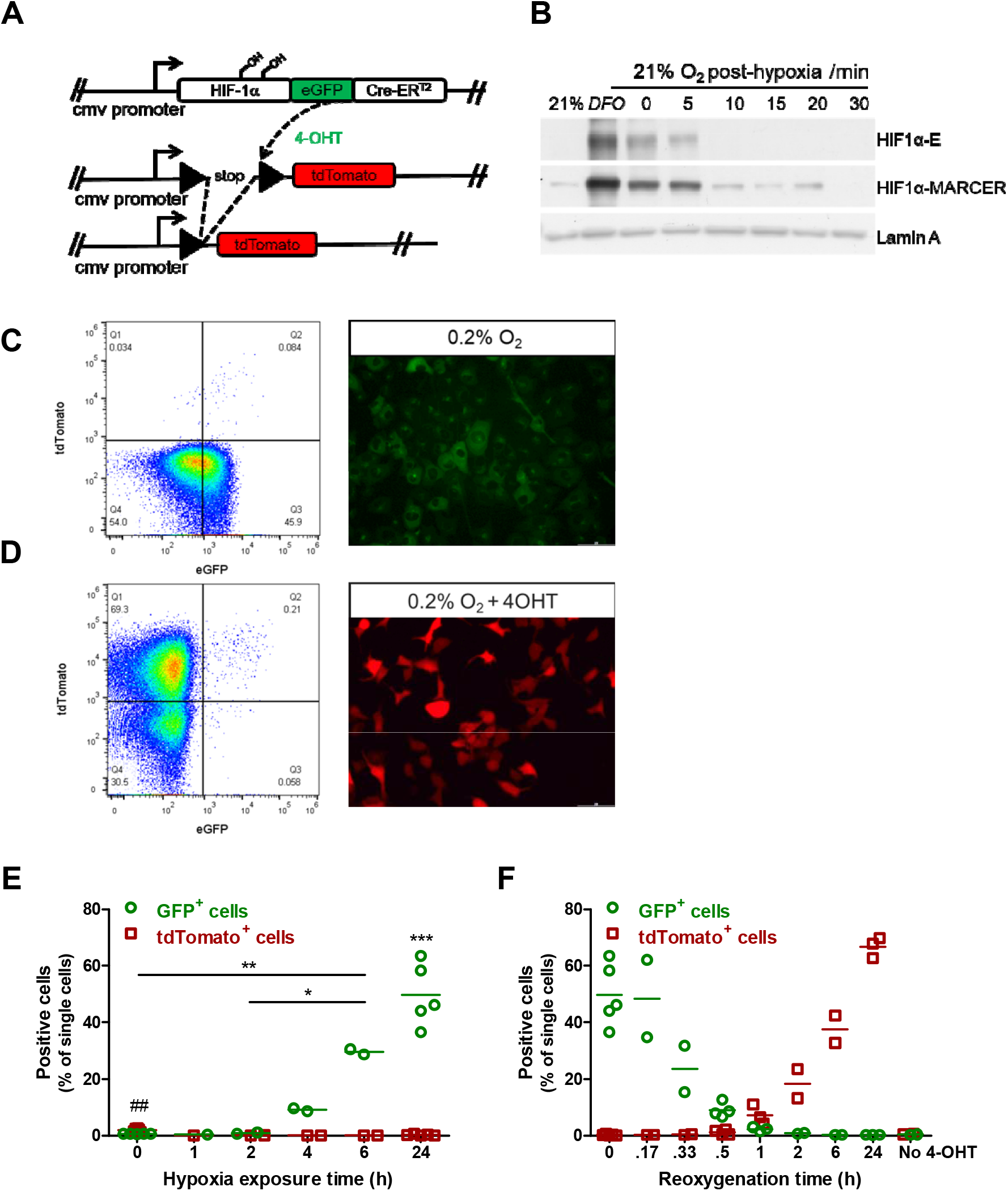
H1299-MARCer reporter (H1299-MR) cells were created. (A) Constructs used for transduction of H1299 cells. (B) Western blot analysis of HIF-1α-Marcer, and endogenous [E] HIF-1α after exposure of H1299-MARCer cells to hypoxia (0.2% O_2_) *in vitro* and re-oxygenation. Lamin A was used a loading control and the HIF-stabilising agent DFO was used as a positive control. (C,D) FACS plots of eGFP and tdTomato expression after exposure to hypoxia (0.1% O_2_) for 24h (C, left) and 24h of hypoxia followed by 24h of re-oxygenation (D, left) Representative images of MARCer stabilisation via eGFP taken after 24h exposure to hypoxia (0.2% O_2_) (C) and after a further 24 reoxygenation of tdTomato expression (D) Scale Bars: 200μm. (E) Flow cytometric analysis of eGFP and tdTomato expression after exposure to increasing times of hypoxia (0.1% O_2_) in the presence of 4-hydroxytamoxifen (4-OHT). Time point ‘0’ is showing cells cultured under normoxia in the presence of 4-OHT for 24h. Dots represent independent experiments carried out in duplicate and coloured bars indicate averages. ##p<0.01 indicates a difference in tdTomato expression between no hypoxia and 2, and p<0.001 for 0 versus 4, 6, and 24h. *p<0.05, **p<0.01 show a difference for eGFP expression as indicated and ***p<0.001 indicates a significantly higher eGFP expression after 24h compared to 0, 2, and 4h, as calculated by One-way ANOVA followed by Bonferroni’s multiple comparison. (F) Flow cytometric analysis of eGFP and tdTomato expression after exposure to 24h of hypoxia (0.1% O_2_) followed by increasing times of re-oxygenation. It should be noted that time point ‘0’ is showing the same data presented in (E) as 24h. Dots represent independent experiments carried out in duplicate and coloured bars indicate averages.

H1299-MR cells were also cultured under hypoxia in the presence of 4-OHT (Fig. 1D, S1B), and eGFP and tdTomato expression were measured by flow cytometry (Fig. 1C-F). Hypoxia (0.1% O_2_) induced the expression of eGFP from 6 hours of treatment onwards (Figs. S1A, 1C,E). MARCer stabilisation is visualised through eGFP expression after 24h exposure to hypoxia (0.2%, Fig. 1C) and cytoplasmic distribution of eGFP after treatment with DFO is visualised in Fig. S1D. tdTomato expression was not visible immediately after exposure to hypoxia and was therefore assessed after reoxygenation for up to 24h and until then tdTomato expression kept increasing while eGFP was rapidly decreased upon reoxygenation (Fig. 1D, F Fig. S1B). In the absence of 4-OHT, tdTomato expression was not induced (Figs.S1C,1F). The HIF1 target gene VEGF was induced by 0,2 % Hypoxia and 4-OHT only slightly further induced these levels (Fig. S1E).

Once tdTomato expression was induced, it was stably expressed in H1299-MR cells under normoxic conditions in the absence of 4-OHT for up to at least 4 weeks (Fig. S1F). When tdTomato^+^ cells were re-exposed to hypoxia, tdTomato expression remained stable (Fig. S1G), however the fluorescence intensity gradually and significantly declined over time (Fig. S1H).

We conclude the HIF MARCer allele reliably reports on endogenous hypoxia and HIF activity, and only slightly increases the HIF transcriptional response when 4OHT is present. By stably inducing tdTomato expression upon administration of tamoxifen we created a reliable tracer of cells exposed to hypoxia with little background fluorescence.

### A single administration of tamoxifen induces tdTomato expression in H1299-MR xenografts

H1299-MR cells were injected subcutaneously into the flank of female Balb/c Nude mice to grow as xenografts. Once tumour size reached approximately 100 mm^3^, tamoxifen was administered by oral gavage and eGFP and tdTomato expression were assessed by flow cytometry 2 days later (Fig. 2A). eGFP expression could not be detected as this is rapidly degraded after exposure to oxygen (Fig. 1B,F) during sample processing. tdTomato was induced by both 5 and 10 mg tamoxifen and 10 mg was used in further experiments as expression appeared more robust (Fig. 2A). tdTomato expression was followed over time and significantly induced from 5 days after tamoxifen administration onwards and expression did not significantly increase beyond 5 days (Fig. 2B).

**Figure 2.**
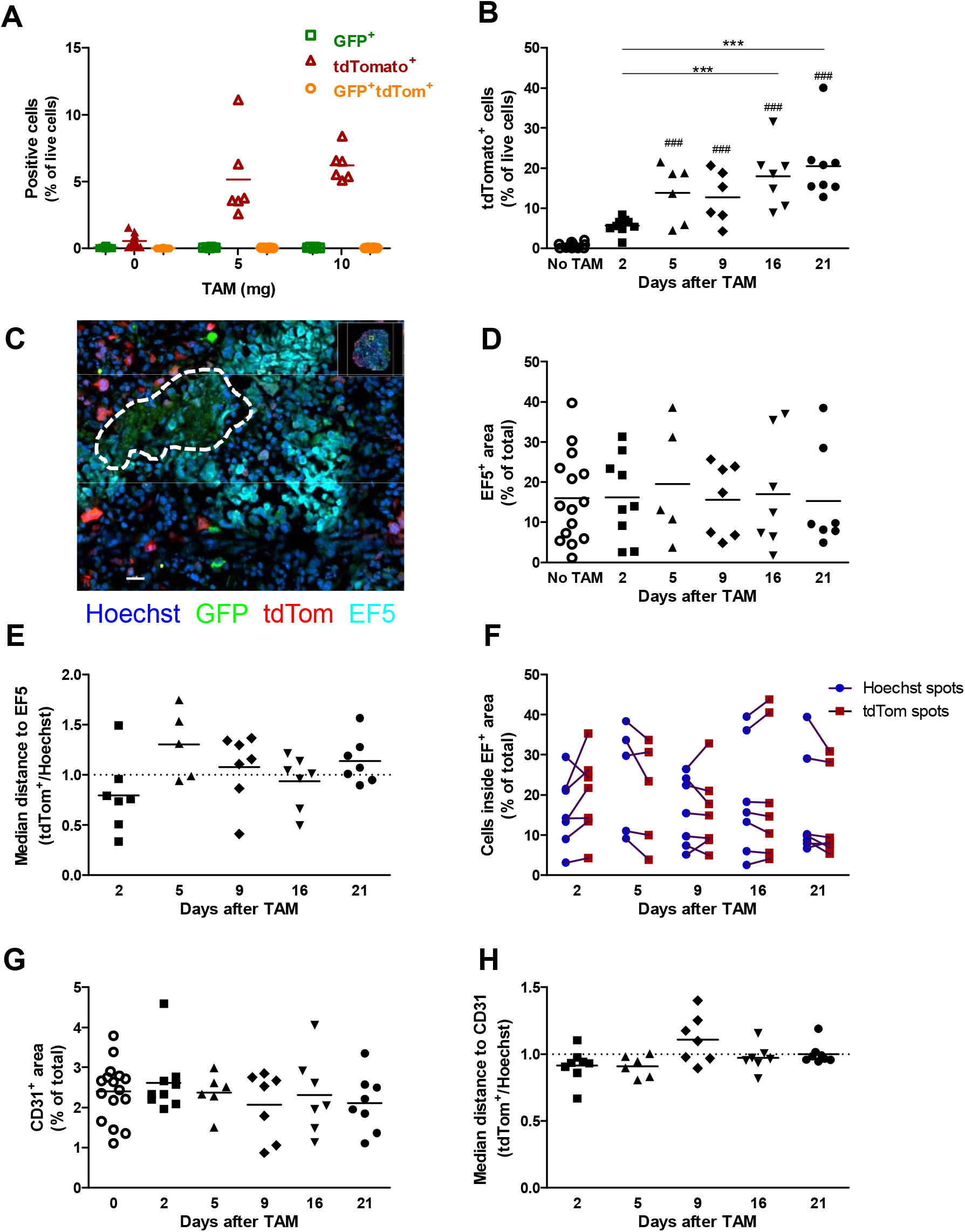
eGFP and tdTomato expression and quantification of immunofluorescent staining of H1299-MR xenografts. (A) One single administration of tamoxifen by oral gavage induced tdTomato expression in H1299-MR xenografts as measured by flow cytometry 2 days after administration. Dots represent individual mice and bars indicate averages (B) From 5 days after administration of 10 mg tamoxifen, tdTomato expression was significantly induced. ###p<0.001 compared to no tamoxifen, ***p<0.001 as determined by One-way ANOVA and Bonferroni’s multiple comparison. (C) Micrograph of EF5 staining showing a necrotic area (dashed line) surrounded by close and more distant EF5-positive staining. Scale bar: 30 μm. (D) EF5 quantification showing hypoxic area as a percentage of total tumour area. (E) Distance of tdTomato-positive cells to EF5-positive areas normalised to all cells (Hoechst, Fig.S2C). (F) Cells inside hypoxic areas as a percentage of all cells or tdTomato-positive cells. (G) CD31 staining showing vessel density as a percentage of total tumour area. (H) Distance of tdTomato-positive cells to CD31-positive areas normalised to all cells (Hoechst, Fig.S2D). For staining (D-H) 1-5 sections per tumour, separated by approximately 1 mm, were analysed and the average was displayed with the dots, whereas the bars indicate the average of all mice in the group.

### tdTomato expression does not significantly correlate with severe hypoxia

To assess whether expression of tdTomato correlated with the extent of hypoxia and vascularisation, we stained frozen tumor sections for EF5 (Figs. 2C, S2A) and CD31 (Fig. S2B). The EF5-positive area did not change over time (Fig. 2D) and did not significantly correlate with tumour size as assessed by a Pearson correlation test on all time points combined (Fig. S3A). EF5^+^ areas were located both in proximity to and more distant from necrotic areas (Figs.2C, S2A). The distance of tdTomato-positive cells to EF5-positive regions was not different from the general population, nor did it change over time (Figs.2E, S2C). tdTomato-positive cells were also equally likely to be inside an EF5^+^ area as the total cell population (Fig.2F). From these results we conclude that post-hypoxic cells are not more likely to reside in hypoxic areas than other tumour cells. Also, the number of tdTomato-positive cells did not correlate with the EF5^+^ area for any of the assessed time points (Fig. S3B). Finally, eGFP-positive cells were occasionally visible (Figs.2C, S2A,B). Quantification of eGFP expression appeared impossible due to high autofluorescence of necrotic areas (Fig.2C, dashed line) and staining for GFP protein did not clearly improve eGFP^+^ cell detectability (not shown).

Tumour sections were stained for CD31 (Fig. S2B) and the percentage of the tumour area covered by vessels was determined. The percentage of vessel area did not change over time (Fig. 2G), nor did the closest distance of each of the tdTomato^+^ cells or of all cells to the nearest vessel (Figs.2H, S2D). Surprisingly, the total areas of EF5 and CD31 did not significantly correlate (Pearson correlation test on all time points combined, Fig. S3C) as we would expect a larger EF5-positive area to correlate with a lower vessel density and therefore the CD31 area. This could be due to areas of increased oxygen demand, limited perfusion, or vessel leakiness in addition to cycling hypoxia.

### RFP staining and tdTomato fluorescence show a similar expression pattern after administration of tamoxifen

Tumour sections were stained using antibodies to RFP by immunofluorescence in order to detect the tdTomato protein (Fig. S4). More RFP-positive cells were detected by immunofluorescence than by imaging intrinsic tdTomato, indicating not all tdTomato-positive cells may be detected by direct fluorescence (Fig. 3A). However, tdTomato fluorescence and RFP immunofluorescence showed a strong correlation (Fig. 3B) and the same expression pattern after tamoxifen (Fig. 3A). A trend that was also similar to tdTomato detection by flow cytometry (Fig. 3C). With imaging on consecutive sections, tdTomato and RFP positive cells were equally likely to be detected inside or outside EF5-positive areas (Fig. 3D). Therefore, even though not all tdTomato cells may be detected with a certain method, this should not introduce bias with regard to lineage tracing of hypoxic cells.

**Figure 3.**
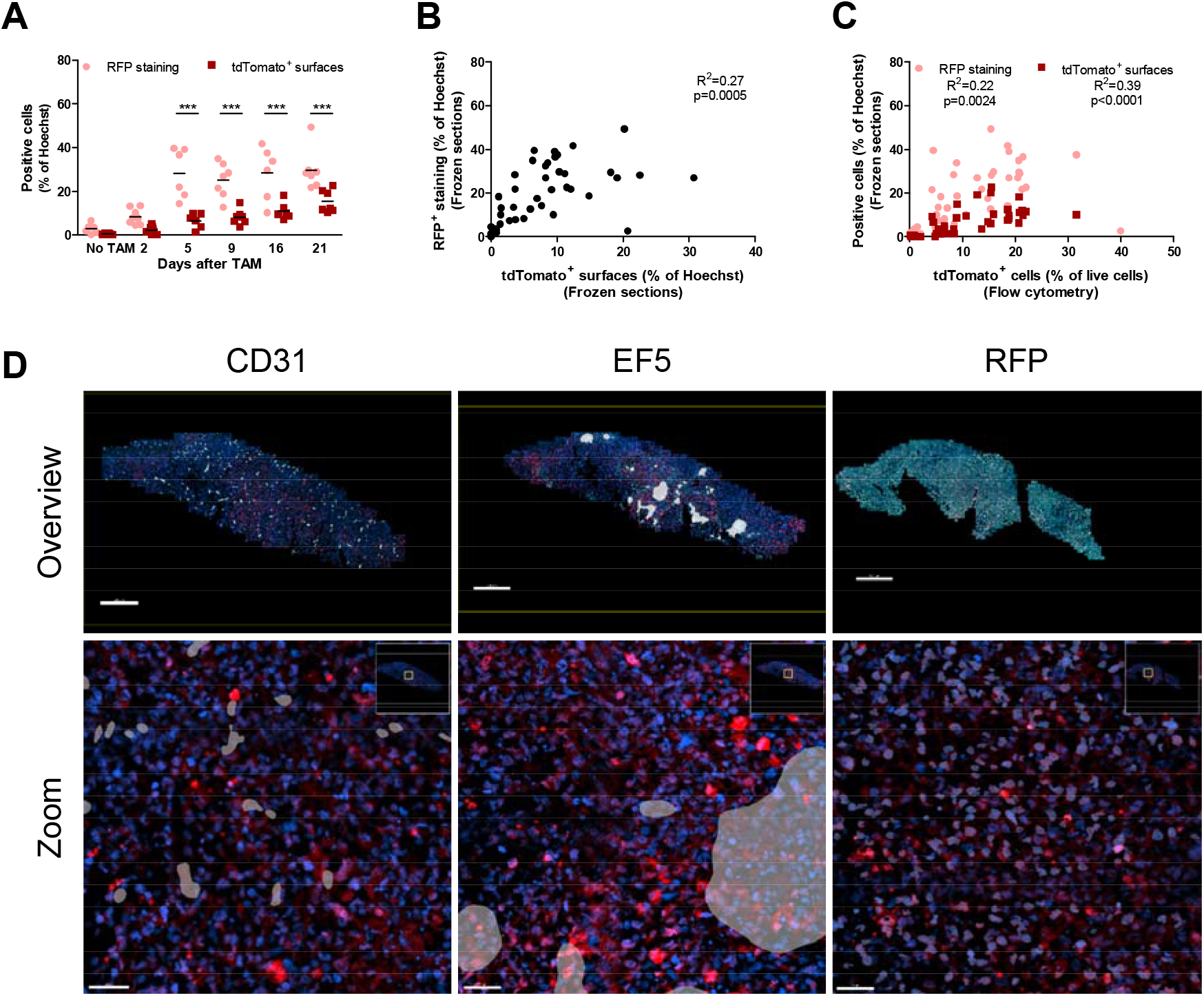
Immunofluorescent staining of H1299-MR xenografts. (A) Quantification of RFP staining showing more tdTomato-positive cells can be detected after staining than when only intrinsic tdTomato was imaged by epifluorescence microscopy. Dots represent the average of 1-5 sections per tumour, separated by approximately 1 mm. ***p<0.001 as determined by Two-way ANOVA followed by Bonferroni’s multiple comparison. (B) RFP staining and intrinsic tdTomato on frozen sections significantly correlate. (C) RFP staining and intrinsic tdTomato significantly correlate with tdTomato cells measured by flow cytometry. (D) Micrographs of consecutive sections showing CD31 staining (left) and RFP staining (right) do not show a clear correlation with EF5 staining (middle). Scale bars top: 500 μm, bottom: 50 μm.

### Post-hypoxic H1299-MR cells proliferate faster than non-hypoxic tumour cells

Next, we assessed the fate of the tdTomato+ post-hypoxic tumour cells. EdU was administered to mice 3 hours before sacrifice and proliferation in xenografts was measured by flow cytometric analysis of EdU incorporation. At all measured time points, tdTomato-positive cells proliferated faster than negative cells comprising both non-hypoxic tumour cells and host cells within the tumour microenvironment (Fig. 4A). This was confirmed by immunofluorescent staining of EdU on 4% PFA-fixed frozen sections (Figs.S5A,4B). Supplementary figure S5B shows that EdU background staining is not significantly higher in tdTomato cells than in all cells. When tumour cells were gated separately from host cells using a human-nuclei-specific antibody (Fig. 4C), tdTomato^+^ cells also proliferated faster than tdTomato^-^ cells (Fig. 4D). These results indicate that tumour cells that were exposed to hypoxia proliferate faster than tumour cells that were not.

**Figure 4.**
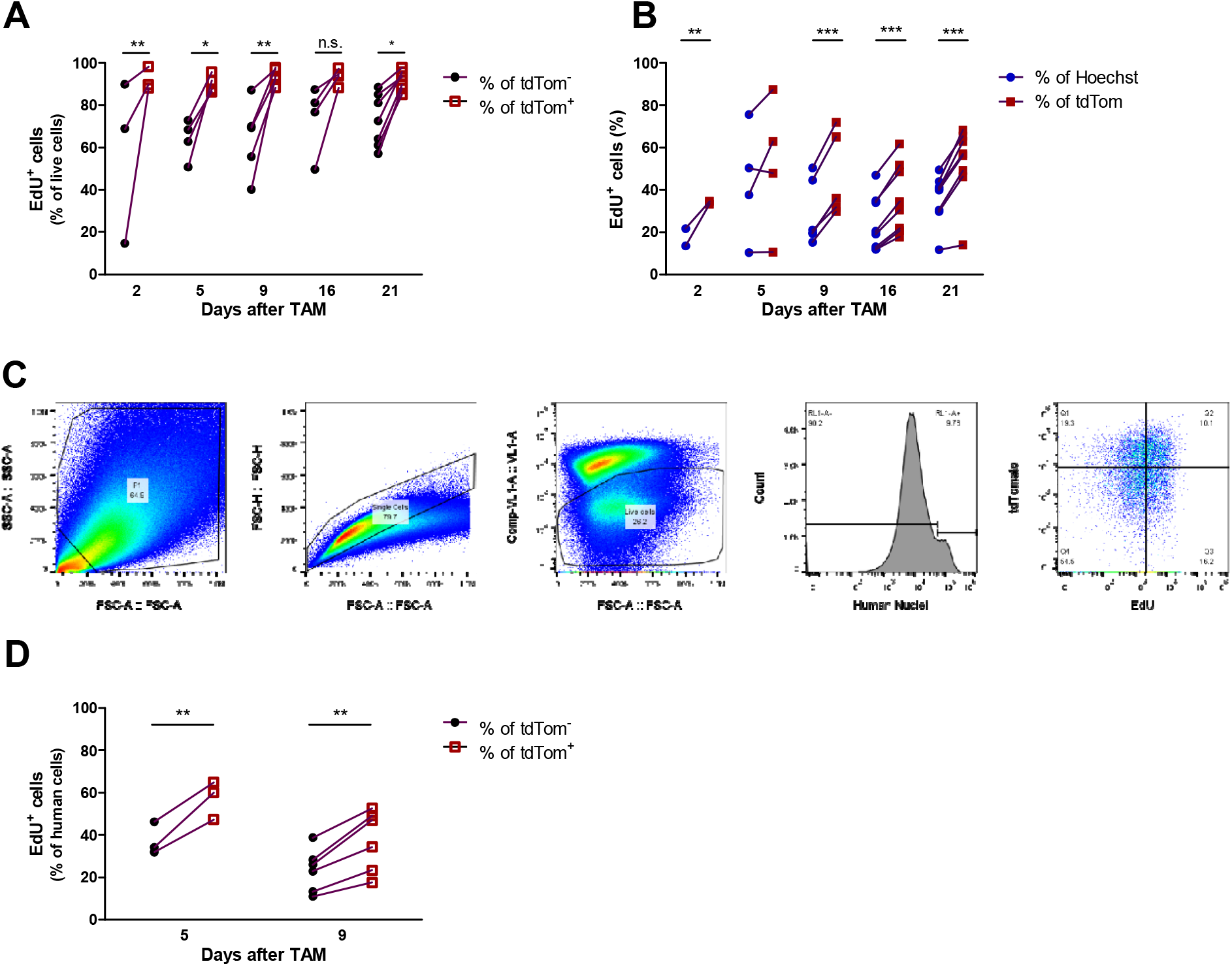
EdU assay showing tdTomato-positive cells proliferated faster than the tdTomato^-^ population as measured by (A) flow cytometry and (B) immunofluorescence. (C) Gating strategy of EdU incorporation in tdTomato^-^ and tdTomato^+^ human cells (RL1-A^+^). (D) EdU proliferation assay showing tdTomato^+^ human cells proliferated faster than the tdTomato^-^ human cells. Dots represent individual mice and paired observations were connected with a line. *p<0.05, **p<0.01, ***p<0.001 as determined by Two-way ANOVA followed by Bonferroni’s multiple comparison.

### Increased proliferation of post-hypoxic H1299-MR tumour cells is non-cell autonomous feature

To address if the increased proliferation was a cell autonomous acquired and stable feature of post-hypoxic tumour cells, H1299-MR xenografts were excised at 5-17 days after administration of tamoxifen and single cell suspensions were cultured *ex vivo.* Antibiotic selection enriched for tumour cells and the percentage of tdTomato-positive cells in culture increased initially after which it stabilised (Fig. 5A). EdU incorporation showed that tdTomato-positive and negative cells proliferated at the same rate at all passages (Fig. 5B). H1299-MR cells were also incubated at 21% and 0.2% O_2_ with 200nm 4-OHT for 24h and mixed in a 1:1 ratio and grown in DMEM containing 1% FBS. Approximately 20-30% of the cells was tdTomato positive and expression did not change as measured by flow cytometry after 3 and 15 days, indicating that proliferation of tdTomato-positive and tdTomato-negative cells was similar *in vitro* (Fig. 5C). These results show that tumour cells previously exposed to hypoxia *in vivo* do not proliferate faster *ex vivo* and that the observed increased proliferation of post-hypoxic tumour cells is influenced by factors in the tumour micro-environment.

**Figure 5.**
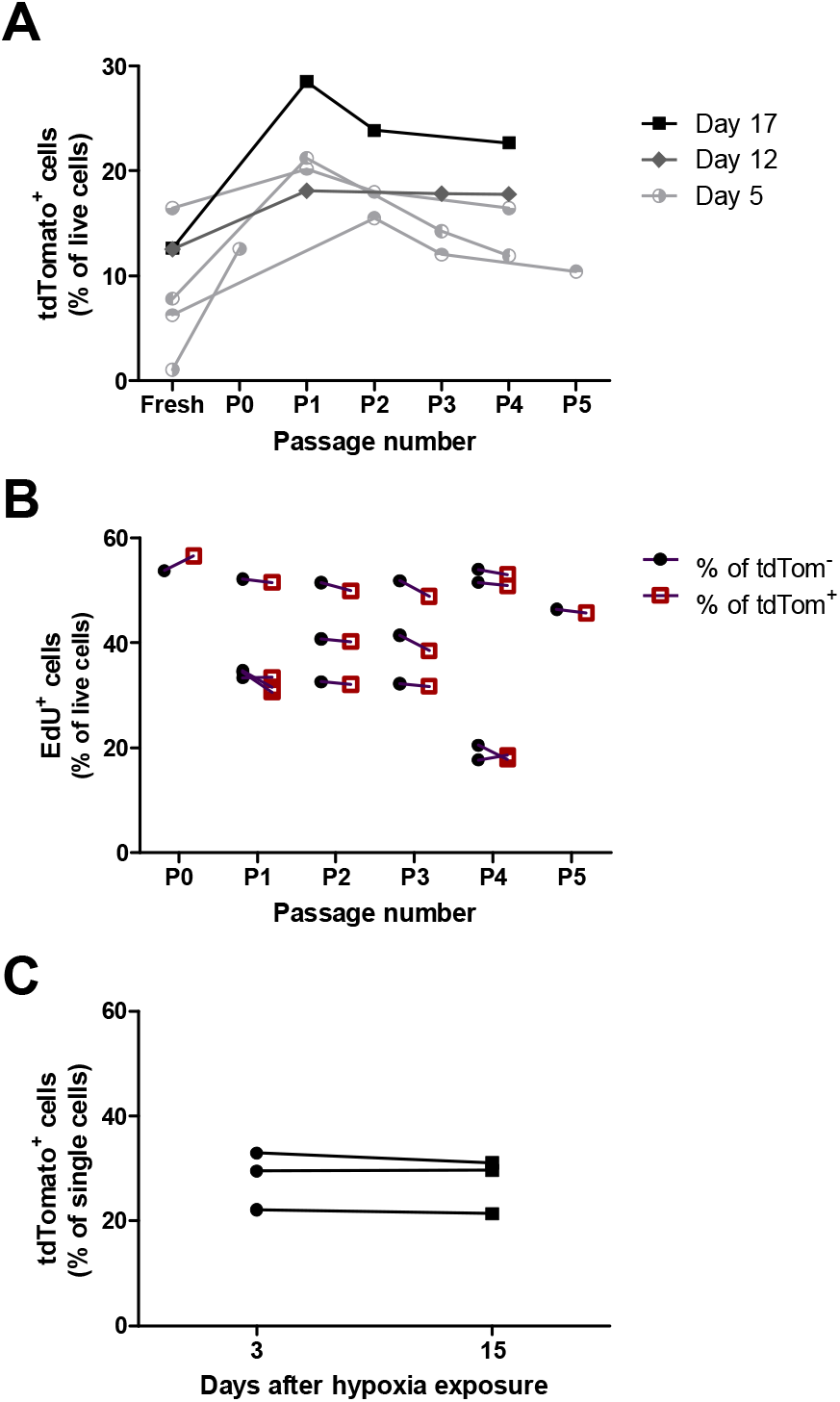
Post-hypoxic tumour cells and non-hypoxic cells proliferate at similar rates *ex vivo* and *in vitro*. (A-B) Cells isolated from H1299-MR xenografts were cultured *ex vivo* under geneticin and blasticidin selection. Individual tumours were depicted and connected with a line. (A) tdTomato expression initially increased and stabilised after several cultured passages as measured by flow cytometry. (B) In *ex vivo* culture, tdTomato^+^ and tdTomato^-^ cells proliferate similarly as shown by EdU incorporation measured by flow cytometry. (C) H1299-MR cells were incubated at 21% and 0.2% O_2_ with 200nm 4-OHT for 24h before being mixed in a 1:1 ratio and grown in DMEM containing 1% FBS. tdTomato expression was then analysed and proved similar after 3 and 15 days.

### Intravital imaging visualises hypoxic cell tracing at the single cell level in xenografts

Next we used intravital microscopy to identify hypoxia lineage tracing *in vivo* in xenograft tumours. H1299-MR xenografts were covered by an imaging window to allow intravital imaging by multiphoton microscopy and tumours were followed for up to 18 days after administration of tamoxifen (Fig. S6). FITC-Dextran or qTracker 705 were injected intravenously and tumours were imaged for vessel perfusion, eGFP, tdTomato, and by second harmonic generation microscopy (Figs. 6A, S6A-D). Fig. 6A is showing an example of a tumour imaged 4 days after administration of tamoxifen and other time points are shown in Fig. S6D. tdTomato was not observed before administration of tamoxifen (Fig. S6B-D). Occasionally eGFP-positive cells were observed (Fig. 6A, arrows). These results demonstrate that we were able to track post-hypoxic tumour cells in a spatiotemporal manner using intravital microscopy.

**Figure 6.**
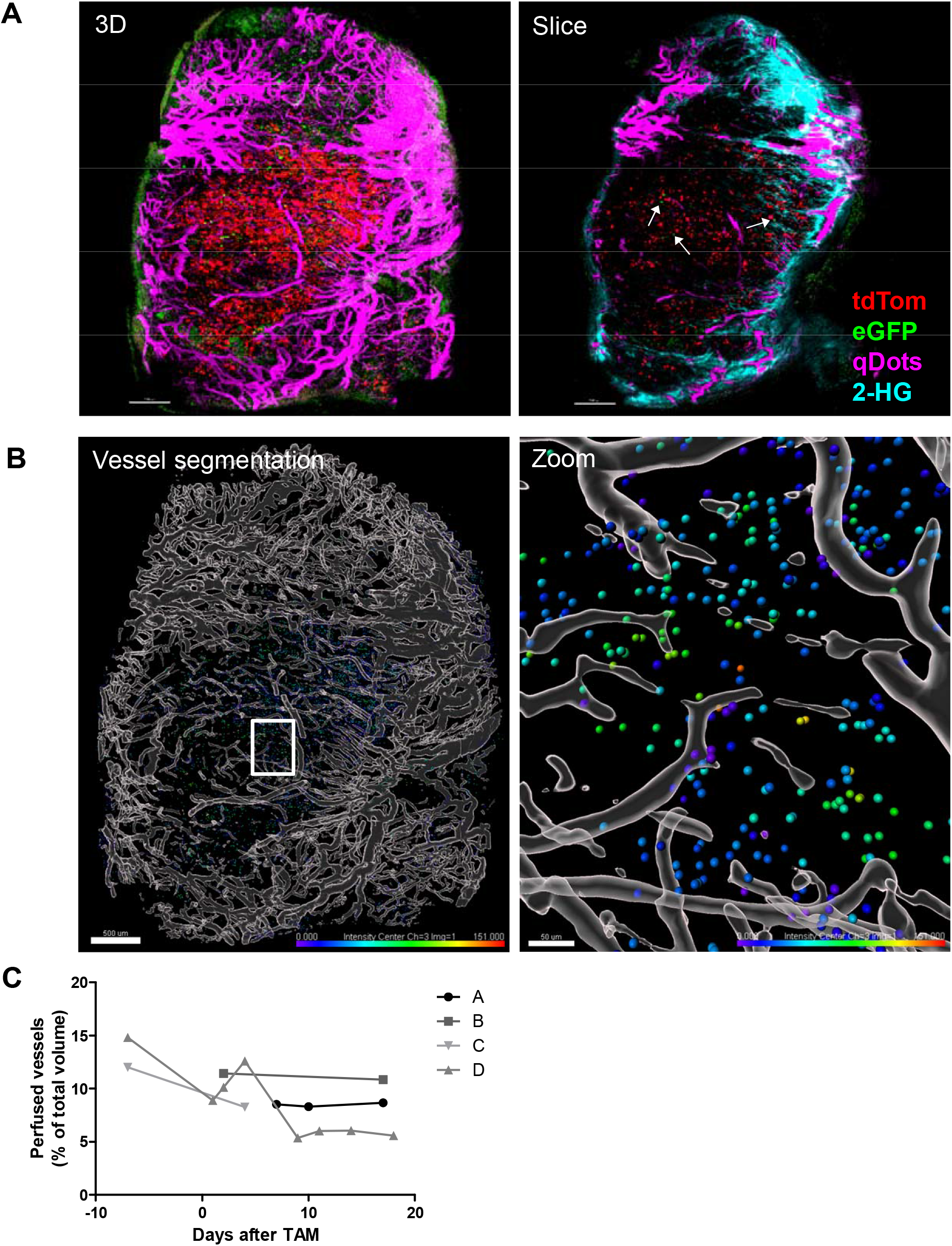
H1299-MR xenografts visualized using intravital microscopy. (A) 3D maximum intensity projection (left) and one slice of the same tumour (right) imaged 4 days after administration of tamoxifen. Perfused vessels are shown in purple, eGFP-positive cells in green, tdTomato-positive cells in red, and collagen in cyan (second harmonic generation microscopy, right panel only). Channel arithmetics was applied using MATLAB to subtract GFP bleed through into the tdTomato channel and tdTomato bleed through into the Qtracker 705 channel. Bars: 500 μm (B) Vessel segmentation (grey) and tdTomato^+^ cells shown in the colour spectrum indicating the distance to perfused vessels (0 μm in purple to 151 μm in red). Bars: 500 μm (left) and 50 μm (zoom). (C) Perfused vessel volume as a percentage of total tumour volume in 4 mice followed over time.

To asses tumour perfusion, we injected qTracker 705 and segmented and reconstructed the vessels with Imaris (Fig.6B). The colour spectrum indicates the distance of tdTomato-positive cells to perfused vessels). The median distance of tdTomato^+^ spots in this tumour was 27.4 μm with a distribution of 0-151 μm. These results indicate that tdTomato cells were located in proximity, as well as further away from the vessels (Fig.6B, right panel), however this was not normally distributed. Cells were more likely to be close to a vessel with 50% of the cells being within 27.4 μm of a vessel in the displayed tumour and an average of 35.5 μm for all tumours (not shown). Purple cells (right panel) with a distance of 0 μm from a vessel appeared to be touching the vessel, rather than circulating inside it. Total vessel perfusion was calculated from the segmented vessels and remained constant over time in 2 of the 4 mice (Fig. 6C, curve A and B, corresponding to mice represented in Fig. S6A-B), whereas vessel volume decreased in mouse C and D (Fig. S6C-D). Overall, tumours seemed well-perfused which is in line with total vessel area as shown by CD31 staining (Fig. 2G).

## Discussion

Here we describe a novel system to lineage trace hypoxic cells. We show that the system is robust *in vitro* with hypoxia-regulated eGFP expression and constitutive tdTomato labelling upon addition of tamoxifen in the progeny of these hypoxic tumour cells. We also see that the MARCer-reporter system does not interfere with endogenous HIF-1α protein expression and the transcription of the HIF target gene VEGF, and this expression is only mildly affected by tamoxifen and only during the short period of tamoxifen administration. *In vivo*, tdTomato expression was induced, however, the amount of expression or the distribution did not correlate with EF5 at any of the investigated time points. Intravital imaging showed that the tumours were well perfused, which is supported by the regular distribution of vessels throughout the tumour and the lack of correlation between vessels and EF5-positive areas as shown by immunofluorescence suggesting also alternative mechanisms of HIF stabilisation.

With the H1299-MR model, *in vitro* exposure to hypoxia induced eGFP expression. However, eGFP was barely detectable *in vivo* with our microscopy systems However, given the clear presence of hypoxic areas as shown by EF5 staining, lack of HIF-1α, and thus eGFP, seems unlikely. This is likely due to the requirement of O_2_ for GFP folding and fluorescence (Heim *et al.*, 1994; Kumagai *et al.*, 2013), making it a suboptimal fluorescent marker to track current hypoxia.. On the other hand, Godet et al. were able to visualise GFP under 0.5% O_2_ (Godet *et al.*, 2019). Also, processing of the samples exposes them to atmospheric oxygen, possibly introducing enough oxygen for GFP maturation *ex vivo*. Despite these limitations we were able to detect eGFP in H1299-MR xenografts using immunofluorescence and intravital microscopy.

Whether limited eGFP expression in our model is related to lack of expression or lack of visibility requires further investigation. An alternative approach could be to replace eGFP with fluorescent proteins not requiring oxygen such as UnaG (Kumagai *et al.*, 2013). Possibly, eGFP expression is truly low in the currently studied tissues, which could be due to the length of time the cells are exposed to cycling hypoxia, not allowing enough time for sufficient MARCer accumulation before reoxygenation and MARCer degradation. This is supported by our current finding that the amount of

EF5 expressed in the tissue does not correlate with the number of tdTomato-positive cells found in the tissue. An alternative explanation is that H1299 NSCLC cells express high levels of reactive oxygen species (ROS), a potent activator of HIF-1α (Jung *et al.*, 2008; Lee, Hwang and Kenyon, 2010). Activation of HIF via ROS could partially explain the lack of overlap and correlation between EF5, HIF-eGFP, and tdTomato labelling.

Surprisingly, neither the fraction of tdTomato^+^ cells and extent of EF5, nor the distance of those cells to EF5 correlated significantly, indicating that post-hypoxic tumour cells are randomly distributed with regard to the current hypoxic status of the tumour. This may be due to the dynamic nature of hypoxia in tumours and the time of assessment, i.e. the time after which the labelled cells were exposed to hypoxia, at least 48h after tamoxifen administration. Because the MARCer construct is constitutively expressed, short episodes of hypoxia could induce labelling, but this may not be seen in proximity to EF5 due to re-oxygenation between the time tamoxifen was given and injection of EF5. Other research has also shown that hypoxic cells move out of hypoxic EF5^+^ or pimonidazole^+^ areas (Harada *et al.*, 2012; Erapaneedi *et al.*, 2016; Conway *et al.*, 2018; Godet *et al.*, 2019) followed by random distribution. The transient nature of the hypoxic state may explain that the induction of tdTomato expression does not correlate with the level of acute hypoxia 2, 5, 9, 16, 21 days later. One way to further investigate this would be through the use of a second marker of hypoxia which is given together with tamoxifen. This would allow visualisation of the hypoxia dynamics and see the change in hypoxia between labelling and endpoint. Taken together EF5 reports on chronic severe hypoxia and anoxia whereas HIF labelling captures also transient tumour hypoxia at moderate O_2_ levels. It also seems evident that many cells that have previously been exposed to hypoxia have redistributed to much less hypoxic regions.

In preliminary results with this system it was apparent that a greater proportion of post-hypoxic cells tdTomato positive cells were undergoing proliferation compared to the non-hypoxic cells. In a range of common cancer types, HIF-1α was positively associated with the proliferation marker Ki67 (Zhong *et al.*, 1999) and several studies report that HIF overexpression in cells can promote cell proliferation (Medici and Olsen, 2012).

While many cell types including cancer cells proliferate more slowly under hypoxia, here we are tracking the uptake of EdU by cells previously exposed to hypoxia. Whether this enhanced proliferation proves to be generalisable remains to be investigated, but might help explain why hypoxia may trigger more aggressive tumours. (Hubbi and Semenza, 2015; Al Tameemi *et al.*, 2019). Blouw and co-workers found a microenvironment-dependent effect of HIF-1α knockdown on tumour progression which might be relevant here. Koshiji et al. (Koshiji *et al.*, 2004) and Hubbi et al. (Hubbi *et al.*, 2013) showed HIF-1α stabilisation induced cell cycle arrest, whereas in our model, HIF-1α was most likely degraded at the time we measured proliferation. It remains to be investigated what mechanisms are responsible for the long-term effect of hypoxia on proliferation in post-hypoxic tumour cells and whether this is dependent on HIF-1α. The H1299-MR model proves to be a promising tool to study this and other long-term effects of hypoxia including its role in metabolic plasticity and metastasis. Moreover, our finding that post-hypoxic tumour cells proliferated faster *in vivo* but not *in vitro* or *ex vivo* demonstrate that this acquired feature is un-stable and non-cell autonomous. It will be of interest to identify what factors in the tumour microenvironment contribute to this.

Other approaches exploiting fluorescent markers driven by hypoxia-responsive elements and oxygen-dependent degradation domains have been described (Erapaneedi *et al.*, 2016; Wang *et al.*, 2016).

Using intravital microscopy, Wang et al. visualised migration of individual normoxic and hypoxic MDA-MB-231 cells in a xenograft model. Similar to our findings, they also reported the presence of hypoxic cells both in proximity and distant from blood vessels. However, they did not quantify whether the distance was different from normoxic cells and different to our current study, Wang et al. studied cells currently experiencing hypoxia whereas we mainly focused on post-hypoxic cells (Wang *et al.*, 2016). Tracing of recently re-oxygenated cells was also performed by Erapaneedi et al. (Erapaneedi *et al.*, 2016), however their system using the fluorescent marker mOrange is dependent on PESTsequence-dependent decay, making it less robust and more challenging to visualise by intravital microscopy, whereas we show stable expression of tdTomato for at least 4 weeks in post-hypoxic cells. Thus, the HIF-MARCer system is a useful addition to the armamentarium to visualize hypoxic, HIF-expressing cells. We show here that H1299-MR cells are a valuable tool to study the long-term fate of hypoxic cells, for example by using intravital microscopy.

In conclusion, our results demonstrate that single-cell lineage tracing of post-hypoxic tumour cells using the H1299-MARCer system allows visualisation of their behaviour in living tumours using intravital microscopy. We provide a valuable tool to study the dissemination and treatment response of post-hypoxic tumour cells *in vivo* and *ex vivo* at a single-cell resolution. Using this system, we provide evidence that post-hypoxic tumour cells may have a proliferative advantage over non-hypoxic tumour cells and that this is influenced by the tumour microenvironment.

## Materials and Methods

### Generating the lenti MARCer system

First, an eGFP CreER^T2^ fusion single primer PCR was performed on plasmid pL451-Dll1(GFP-ires-CreER^T2^) (van Es *et al.*, 2012) a kind gift of Johan van Es, with primer: 5’-GGC ATG GAC GAG CTG TAC AAG TCC AAT TTA CTG ACC GTA CAC-3’ and on pEGFP-C1 (Clontech) with primer: 5’-GGC ATG GAC GAG CTG TAC AAG-3’. After the reaction these template plasmids were digested with DpnI, before mixing 1ul of each reaction for a fresh PCR with primer pair 5’-GTG AGC AAG GGC GAG GA-3’ and 5’-CCA GAC ATG ATA AGA TAC ATT GAT GAG-3’ to amplify the fusion product with proofreading Phusion DNA Polymerase (FynnZyme). This GFP-CreER^T2^ PCR fragment was gel purified, T overhangs were added with normal Taq polymerase before sub cloning it in a pCR-XL-TOPO^®^ vector (Invitrogen) generating pCR-GFP-CreER^T2^ vector. Afterwards this vector was used as a template to generate in a similar approach the fusion with HIF-1α. A HIF-1α ~1.5kb gDNA fragment was amplified from BAC clone RP11: 618G20 (GenBank:AL137129.4), using primers 5’-TGG ATC CGA GCT CGG TAC CAT AGA TCT GAA CAT TAT AAC TTG ATA AAT GAG G-3’ and 5’-AGC TCC TCG CCC TTG CTC ACC TGG AAT ACT GTA ACT GTG C-3’. The product, a fusion of HIF-1α in exon 12 with 20nt of the cDNA GFP, was cloned by use of the NEBuilder^®^ HiFi DNA assembly cloning kit in the pCR-GFP-CreER^T2^ plasmid generating pCR-HIF-LHA-GFP-CreER^T2^. On this plasmid a single primer PCR with primer 5’-GCA AGC CCT GAA AGC GCA AG-3’ and the cDNA human HIF-1α in a p3XFLAG-CMV™-10 expression vector (Sigma, St Louis, MO, USA) (Gort *et al.*, 2008) was used as a template with a complementary single primer PCR with primer: 5’-CTT GCG CTT TCA GGG CTT GC-3’. After the reaction the template plasmids were digested with DpnI, before mixing 1ul of each reaction for a fresh PCR with primer pair 5’-GAT ATC GGT ACC AGT CGA CTC-3’ and 5’-GTG GTA CCC GTC ATC AAG CTG TGG CAG GGA-3’ amplifying the MARCer cDNA (Fig.1A, top) which was sub cloned in a pCR^®^-Blunt II-TOPO^®^ vector generating pCR-BluntII-MARCer. Flanked by BstXI sites the MARCer cDNA was, after digestion, retrieved by gel purification to replace luciferase in the BstxI digested pLenti CMV V5-LUC Blast (w567-1), that was a gift from Eric Campeau (Addgene plasmid # 21474) (Campeau *et al.*, 2009), generating pLenti-CMV-MARCer-Blast. This plasmid expresses amino acid 1-603 of the human HIF-1α protein fused to eGFP-CreER^T2^.

The Ai65(RCFL-tdT) targeting vector Addgene Plasmid #61577 which was a gift from Hongkui Zeng (Li *et al.*, 2015) was digested with BstBI-AscI and a 6267bp fragment was isolated and cloned into an empty BstBI-AscI digested pLVX-puro vector with an introduced unique AscI site to fuse proteins to FLAG and HA tags at the carboxy terminus (Groot *et al.*, 2014). Next, the FLAG-HA-PGK-Puro-WPRE fragment was removed from this vector with an AscI-KpnI digest followed by blunting of the DNA ends with Mung Bean and ligated back. Finally, the FRT-STOP-FRT cassette was removed with a digest with XhoI and the vector was back ligated to generate the pLV(cmv)-NEO-SFS-tdTomato Cre-reporter plasmid. The full integrity of all constructs was confirmed by DNA sequencing.

### Generation of the H1299-MR cell line

Viral particles were produced using viral vectors and packaging plasmids in 293FT cells as previously published (Groot *et al.*, 2014). H1299 on which we performed STR analysis were first transduced with MARCer viruses and cells were selected with 10 μg mL^-1^ of Blasticidin in the 10% FCS RPMI culture medium, supplemented with Penicillin/Streptomycin. Transduced cells were single cell seeded to form clones on 15 cm dishes. Next, clones were harvested by use of glass clonal cylinders (Sigma-Aldrich) with Baysilone-Paste (GE Bayer Silicones) and expanded. Clones were split and transiently transfected with the reporter plasmid described above and screened for their switching capacity with the addition of DFO 100 nM and 100 nM 4-hydroxytamoxifen (H7904-5MG, Sigma-Aldrich). We identified clone number 12 to show most tdTomato expression after treatment. Next, clone 12 was subsequently transduced with SFS-tdTomato viruses and clones were selected after this second transduction and were selected with 10 μg mL^-1^ of Blasticidin and 1000 μg mL^-1^ G418 in the 10% FCS RPMI culture medium, supplemented with Penicillin/Streptomycin. The polyclonal cell population was exposed to 1% of oxygen for 24 h in a Russkinn INVIVO 1000 hypoxic chamber. Next the recovered cells were passed in culture at normoxic conditions for two days. To these cells 100 nM of 4-OHT was added to the medium for 24 h. To select for cells without leakage, cells were single cell seeded and colourless clones were picked as described above. Clones were expanded and split in 250k cells per 6 well and exposed to 200 nM 4-OHT under hypoxia and normoxia for 24 h, trypsinised and expanded for three days in culture flasks. We identified H1299 clone 12.3 (H1299-MR) as a robust hypoxia reporter cell line that we characterised and used in our further experiments.

### Cell culture and hypoxia

H1299-MR cells were cultured in 10% fetal bovine serum (FBS, Gibco) RPMI culture medium (Sigma-Aldrich) supplemented with 1% Penicillin/Streptomycin (Sigma-Aldrich), 10 μg mL^-1^ of Blasticidin (Invivogen) and 1000 μg mL^-1^ G418 (Sigma-Aldrich). Cells were regularly tested for mycoplasma and incubated in a humidified incubator or hypoxia chamber Ruskinn InVivo 300 (Figs.1C&D left panels, E-F, S1A,B,G-H) or InVivo 1000 (Figs.1B, 1C-D right panels, S1C-F, 5C) with or without 4-OHT. Images during cell culture were taken with a Nikon eclipse Ts2 microscope.

### Western blotting

Cells were lysed in RIPA buffer, containing 50 mM Tris·HCl, 0.5% DOC, 0.1% SDS, 1 mM EGTA, 2 mM EDTA, 10% glycerol, 1% Triton X100, 150 mM NaCl, and 1 mM PMSF, and protease inhibitors (ROCHE pill Complete Inhibitor). Protein concentrations were determined with Bradford Protein Assay (Bio-Rad). Proteins (30 g) were separated on a 6% SDS-PAGE and transferred to a PVDF membrane. The membrane was blocked with 5% non-fat dry milk (Marvel) and subsequently incubated (O/N, 4°C) with primary antibodies HIF-1α (BD Biosciences) Lamin A (Sigma) and then horseradish peroxidase-linked secondary antibodies (horse anti-mouse and horse anti-rabbit; 1:2,500; Cell Signaling). ECL Prime Western Blotting Detection Reagent (GE Healthcare) was used for visualisation.

### RNA isolation and qPCR

mRNA was extracted using the Nucleospin RNA II kit (Bioke), and cDNA conversion was performed using iScript cDNA Synthesis kit (Bio-Rad) according to the manufacturer’s instructions. Quantitative PCR was performed on the CFX96 (Bio-Rad). The expression of VEGF (F: 5’-GACTCCGGCGGAAGCAT-3’ R: 5’-TCCGGGCTCGGTGATTTA-3’) was detected with SYBR Green I (Eurogentec). Gene expression was normalised to RPL13A (F: 5’-CCGGGTTGGCTGGAAGTACC-3’ R: 5’-CTTCTCGGCCTGTTTCCGTAG-3’) mRNA expression.

### Mice

All animal studies were performed according to the Animals Scientific Procedure Act 1986 (UK) and approved by local ethical review. Female balb/c nude mice were obtained from Envigo and kept in individually ventilated cages with unlimited supply of food and water and 12 h light-dark cycles and mice were weighed twice a week. H1299-MR cells were harvested, dissolved in a single-cell suspension with Matrigel (Corning Life Sciences, 1:1) and 10^6^ cells were injected subcutaneously into the flank of 6-to-10-week-old mice. Once tumours reached approximately 100 mm^3^, tamoxifen (TAM, Sigma-Aldrich) was dissolved in vegetable oil containing 5% ethanol and administered through oral gavage. At the end of the experiment, mice were injected intraperitoneally with EF5 (2-(2-nitro-1/-/-imidazol-l-yl)-N-(2,2,3,3~-pentafluoropropyl)acetamide, a nitroaromatic compound stabilized in the absence of oxygen (Lord et al. 1993 PMID: 8242628), a kind gift from Prof. Cameron Koch, University of Pennsylvania) and EdU (5-Ethynyl-2’-deoxyuridine, Santa Cruz) 3 hours before sacrifice. Tumours were harvested, rinsed in PBS, cut in half and processed for flow cytometry or immunofluorescence.

### Flow Cytometry

Tumours were collected, halved, kept in HBSS (Gibco) and chopped into small pieces using a scalpel. Tumours were then digested in HBSS (Gibco) containing Collagenase Type 2 (Worthington Biochemical Corporation) and DNAse I (ThermoFisher Scientific) for 40 min at 37°C in a shaking incubator. Cells were filtered through a 50 μm strainer (Sysmex Partec) and rinsed with FACS Buffer (PBS containing 5% FBS) and centrifuged for 5 min at 300g, 4°C and rinsed again. Cells were incubated for 15 min with LIVE/DEAD Fixable Violet Dead Cell Stain (ThermoFisher Scientific) and a maximum of 2×10^6^ cells was used for EdU staining using the Click-iT EdU Alexa Fluor 488 Flow Cytometry Assay according to manufacturer’s instructions (ThermoFisher Scientific). The remaining cells were fixed using IC fixation buffer (1:1 in PBS, ThermoFisher Scientific) for approximately 10 min at room temperature. Cells were stored in IC fixation buffer at 4°C until analysis on the Attune NxT Flow Cytometer (ThermoFisher Scientific) when they were centrifuged for 5 min at 400g and dissolved in FACS buffer. For compensation purposes, H1299-MR cells were cultured *in vitro* and subjected to DFO treatment (eGFP, channel BL-1), TAM + hypoxia and reoxygenation (tdTomato, channel YL-1), 5 min at 65°C (LIVE/DEAD, channel VL-1, 1:1 with untreated cells), EdU (Click-iT assay, channel BL-1) or left untreated (unstained control and human nuclei, channel RL-1). When stained for human nuclei (MAB1281, Merck Millipore, Fig. 4C-D), this was performed for 30 min at RT in the dark after EdU staining (Click-iT Assay) and blocking. AF647 donkey anti-mouse IgG (ThermoFisher Scientific) was used as a secondary antibody.

Cells cultured *in vitro* that were used for flow cytometry were rinsed and scraped in PBS, and added to IC fixation buffer (1:1 in PBS) inside the hypoxia chamber. They were then taken out of the chamber and incubated, together with cells not exposed to hypoxia, for approximately 30 min at RT. Cells were stored in IC fixation buffer at 4°C until analysed, they were centrifuged for 5 min at 400g and dissolved in FACS buffer. An example of the gating strategy was shown in Fig. 1C-D and analyses were performed in FlowJo (BD).

### *Ex vivo* cell culture

When cells from xenografts were cultured *ex vivo*, tumours were harvested and put in RPMI complete medium without selection antibiotics. They were then digested as described above. From this point, cells were kept under sterile conditions, filtered through a 30 μm strainer (Sysmex Partec) and rinsed with PBS containing 2% BSA and 5 mM EDTA (both Sigma-Aldrich). After centrifugation for 5 min, 300g at 4°C, the pellet was resuspended and incubated for 3 min in red cell lysis buffer (155 mM NH_4_Cl, 12 mM NaHCO_3_, 0.1 mM EDTA). Cells were rinsed twice with PBS/BSA/EDTA and dissolved in PBS. Half of the cells was taken into culture with RPMI complete medium and 10 μg mL^-1^ of Blasticidin and 1000 μg mL^-1^ G418 and the other half was stained for LIVE/DEAD, fixed, and analysed by flow cytometry. At several culture passages, cells were harvested with Trypsin/EDTA (Sigma-Aldrich), filtered through a 30 μm strainer and stained for LIVE/DEAD and EdU as described above, and analysed by flow cytometry.

### Immunofluorescence and Microscopy

Tumour halves were rinsed in PBS and fixed for 3-4 h in 4% PFA at 4°C. They were transferred to a 30% sucrose in PBS solution and kept overnight in the fridge and consecutively snap frozen in OCT embedding medium (ThermoFisher Scientific) and stored at −20°C until cutting. 10 μm thick cryosections were cut using a Bright Cryostat or a Leica CM1950, dried overnight at 37°C and stored at −80°C until staining.

Sections were allowed to dry at room temperature for at least 30 min, rinsed in PBS and permeabilised with 0.5% Triton-X-100 in PBS. For staining of CD31, 5% BSA (Sigma) and 5% donkey serum (Sigma-Aldrich) were added to the permeabilisation solution and this was also used for blocking for 1h at RT. Blocking for RFP staining was done with 10% normal goat serum (Sigma-Aldrich) in PBS for 1h at RT and with TNB blocking buffer (PerkinElmer) for 30 min for EF5 staining. Staining was performed overnight at 4°C with Cy5-conjugated EF5 antibody or Cy5-EF5 antibody containing competitor (both purchased from University of Pennsylvania and diluted 1:1 in PBS), rabbit anti-RFP (600-401-379, Rockland) or rabbit IgG (BD Biosciences) at 1:500 in PBS, and goat anti-CD31 (AF3628, R&D Systems) or goat IgG (R&D Systems) at 1 μg mL^-1^. Sections stained for EF5 were washed with PBS/Tween-20 0.3% twice for 45 min. Sections stained for RFP and CD31 staining were washed 3x in PBS and incubated for 30 min at RT with secondary antibodies AF633 goat anti-rabbit and AF647 donkey anti-goat, respectively. The Click-iT EdU Cell Proliferation Kit for Imaging (ThermoFisher Scientific) was used according to the manufacturer’s instructions and a solution containing only the Alexa Fluor picolyl azide in PBS/BSA 3% was used as a negative control. All sections were counterstained with Hoechst 33342 (ThermoFisher Scientific) and mounted in Prolong Diamond Antifade Mountant (ThermoFisher Scientific). Entire sections were imaged using a Nikon-NiE epifluorescence microscope with a 20x objective, images were stitched with NIS Elements (Nikon), and processed with Imaris software (Bitplane). To get an overview of the entire tumour, 1-5 sections per tumour, separated by approximately 1 mm, were analysed and the average was displayed.

### Abdominal Imaging Window and Intravital Microscopy

Under inhalation anaesthesia with isoflurane, an imaging window was placed onto the abdominal wall of a mouse as previously described (Ritsma *et al.*, 2013) with the following adjustments. A titanium window was used and a coverslip was glued on top with BBRAUN Histoacryl (Akin Global Medical). Vetergesic was injected subcutaneously as analgesic. An incision was made in the skin, after which 5×10^5^ H1299-MR cells in 30% matrigel in 5 μL were injected into the thin fat layer above the abdominal muscle. The window was stitched to the muscle layer using 5-0 silk and then secured to the skin with 5-0 prolene sutures (Ethicon Inc.). After approximately 6 weeks, when tumours became visible by eye, 10 mg tamoxifen in 100 μL 5% ethanol/oil mixture was administered by oral gavage. At every imaging session, windows were carefully cleaned with an insulin syringe using 0.9% NaCl and all liquid and air between the coverslip and the tumour was removed. FITC-Dextran (MW 500, Sigma-Aldrich) or Qtracker 705 (ThermoFisher Scientific), diluted 1:10 in sterile saline, was administered via the cannulated tail vein by a bolus injection of 2×12.5 μL, followed by a rate of 70 μL h^-1^ using an automated pump (Harvard Instruments). Tumours were imaged using a ZEISS LSM 880 microscope with a Mai-Tai laser (Newport Spectra-Physics, 940 nm excitation) using a ZEISS 20x 1.0 NA water objective covered with ultrasound gel. For detection of Qtracker 705, a 670 nm shortpass filter was used, whereas bandpass filters were used of 457-487 for collagen (second harmonic generation microscopy, 2-HG), of 488 to 512 nm for eGFP/FITC, and 562.5 to 587.5 nm for detection of tdTomato. Tile scans were taken up to 350 μm deep with a voxel size of 0.83×0.83 in x-y and 5 μm in z, and stitched using ZEN Black Software (Carl ZEISS AG). Quantitative analyses were performed using Imaris.

### Image Analyses

All analyses were performed using Imaris software. First, the total tumour was outlined, background such as folds were excluded as much as possible and signal outside the tumour area was removed. For EF5 staining, hypoxic areas directly underneath the skin were excluded from the analysis. For stained sections, masks were created for Hoechst, tdTomato, and staining using the ‘surfaces’ or ‘spots’ functions and thresholds adjusted for each imaging session. All analyses were checked by eye and when the mask did not visually represent the positively-stained area, the data were excluded from the analysis. A distance map from CD31^+^ and EF5^+^ areas was created using MATLAB and the median distance of tdTomato^+^ spots and Hoechst^+^ spots to CD31 and EF5 surfaces was calculated.

For analyses of EdU, Hoechst^+^ cell surfaces were masked. These were filtered for an EdU^+^ threshold, either or not preceded by the tdTomato^+^ threshold and expressed as a percentage of total. RFP^+^ cells were also filtered by an intensity threshold on Hoechst-positive surfaces.

For intravital images, channel arithmetics was applied using MATLAB to subtract GFP bleed through into the tdTomato channel and tdTomato bleed through into the Qtracker 705 channel. A surfaces mask was created on the Qtracker 705 signal and the perfused-vessel volume was expressed as a percentage of total tumour volume. A distance map from vessels was created using MATLAB and the distance of tdTomato^+^ spots to vessels was represented as a colour spectrum (0 - 151 μm).

### Statistical Analyses

Statistical analyses were performed using GraphPad Prism software. This was also used to create the figures which are showing the mean and individual measurements carried out in duplicate unless stated otherwise. Paired observations are displayed with a connecting line. The statistical tests performed are indicated for each figure and p<0.05 was considered significant.

## Acknowledgements

JI and AG and LB were supported by the European Research Council under the European Union H2020 research and innovation programme (grant 617060 to MV). The research leading to these results has also received funding from the People Programme (Marie Curie Actions) of the European Union’s Seventh Framework Programme (FP7/2007-2013) under REA grant agreement No 625631 to BM. RJM received funding from Cancer Research UK (grant C5255/A15935). We thank Prof. Cameron Koch at the University of Pennsylvania for providing EF5.

We would like to thank Graham Brown and Rhodri Wilson at the Microscopy Scientific Research Facility for advice regarding microscopy and image analysis, John Prentice for help with creating the imaging windows, and Karla Watson, James Ward, and Magda Hutchins at Biomedical Services for technical support and animal care.

