## Supplementary figures and legends for "A lineage-tracing tool to map the fate of hypoxic tumour cells"

**Figure S1.**

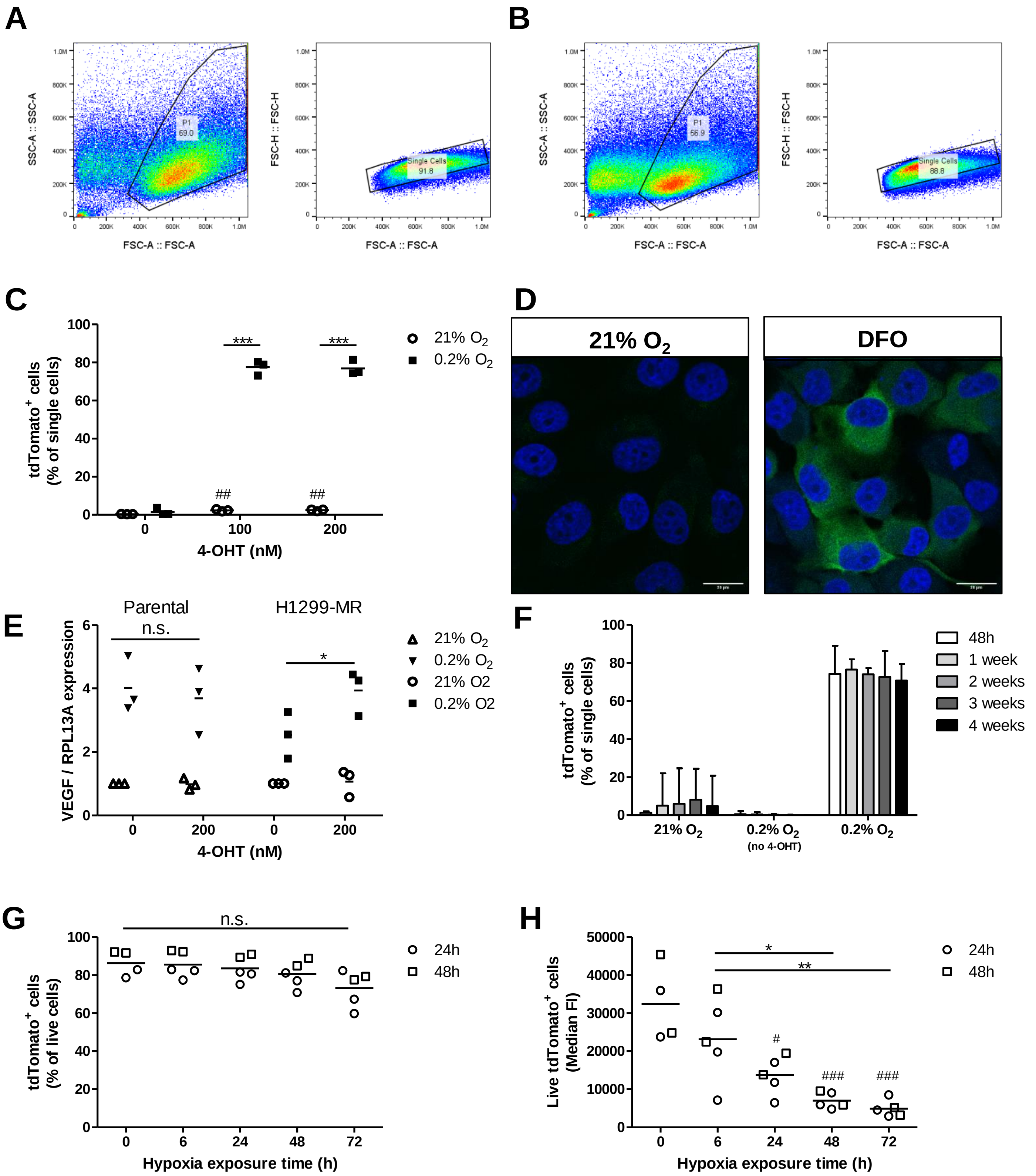

A
B
SSC-A :: SSC-A
FSC-A :: FSC-A
FSC-H :: FSC-H
P1
69.0
Single Cells
91.8
P1
56.9
Single Cells
88.8
C
tdTomato+ cells (% of single cells)
4-OHT (nM)
21% O2
0.2% O2
***
##
D
21% O2
DFO
E
Parental
H1299-MR
n.s.
*
VEGF / RPL13A expression
4-OHT (nM)
21% O2
0.2% O2
21% O2
0.2% O2
F
tdTomato+ cells (% of single cells)
48h
1 week
2 weeks
3 weeks
4 weeks
21% O2
0.2% O2 (no 4-OHT)
0.2% O2
G
n.s.
tdTomato+ cells (% of live cells)
Hypoxia exposure time (h)
24h
48h
H
Live tdTomato+ cells (Median FI)
Hypoxia exposure time (h)
*
**
#
###
24h
48h

**Figure S1. H1299-MR characteristics.** Gating strategy of eGFP and tdTomato expression after exposure to hypoxia (0.1% $O_2$) for 24h (A) and 24h of hypoxia followed by 24h of re-oxygenation (B) (C) Flow cytometric analysis of tdTomato expression 72 hours after 0.2% $O_2$ and different concentrations of 4-OHT. *** $p<0.001$ as indicated using 2-way ANOVA followed by Bonferroni post-test. ## $p<0.01$, 4-OHT versus without 4-OHT under normoxia using 1-way ANOVA followed by Bonferroni's multiple comparison. (D) Distribution of eGFP after 24h treatment of H1299-MR with 100µM DFO. Scale bar: 20µm. (E) VEGF gene expression normalised for the housekeeping gene RPL13A after exposure to tamoxifen in the parental H1299 cell line and H1299-MR Dots represent independent experiments carried out in triplicate and bars indicate averages. (F-H) H1299-MR cells stably expressing tdTomato were re-exposed to hypoxia. (F) Stable expression of tdTomato in normoxia once it was induced by hypoxia (0.2% $O_2$) and 4-OHT Bars represent mean expression of 3 independent experiments and error bars indicate 95% c.i. (G) tdTomato was induced with 4-OHT and 24h (circles) or 48h (squares) of 0.1% $O_2$ as indicated. Re-exposure to hypoxia for up to 72 hours did not significantly affect the percentage of tdTomato$^+$ cells. Dots represent independent experiments carried out in duplicate and bars indicate averages (H) Re-exposure to hypoxia gradually reduced the tdTomato median fluorescence intensity. Dots represent independent experiments carried out in duplicate and bars indicate averages #$p<0.05$, ###$p<0.001$ compared to no hypoxia re-exposure and *$p<0.05$, **$p<0.01$ as indicated and calculated by One-way ANOVA followed by Bonferroni's multiple comparison.

**Figure S2.**

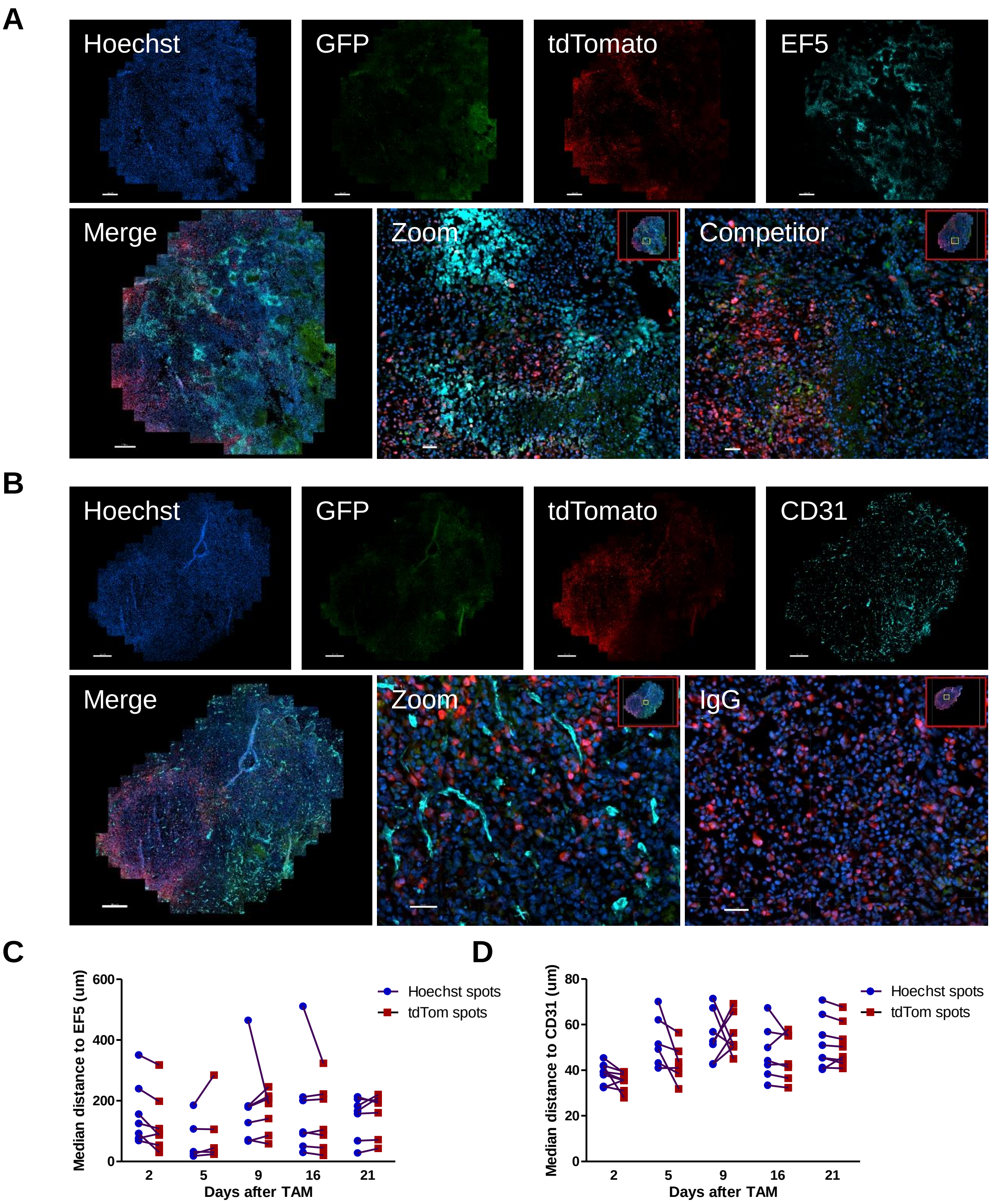


**Figure S2. Immunofluorescent staining of H1299-MR xenografts at different time points after administration of tamoxifen.** (A) Micrographs of EF5 staining. An EF5-Cy5 antibody mixture containing a competitive inhibitor of the antibody was used as a negative control (competitor). (B) CD31 and IgG control staining (C) Distance of tdTomato-positive cells and all cells to EF5-positive areas. (D) Distance of tdTomato-positive cells and all cells to CD31-positive areas. Scale bars overview: 500 µm, scale bars zoom and controls: 50 µm.

### Figure S3.

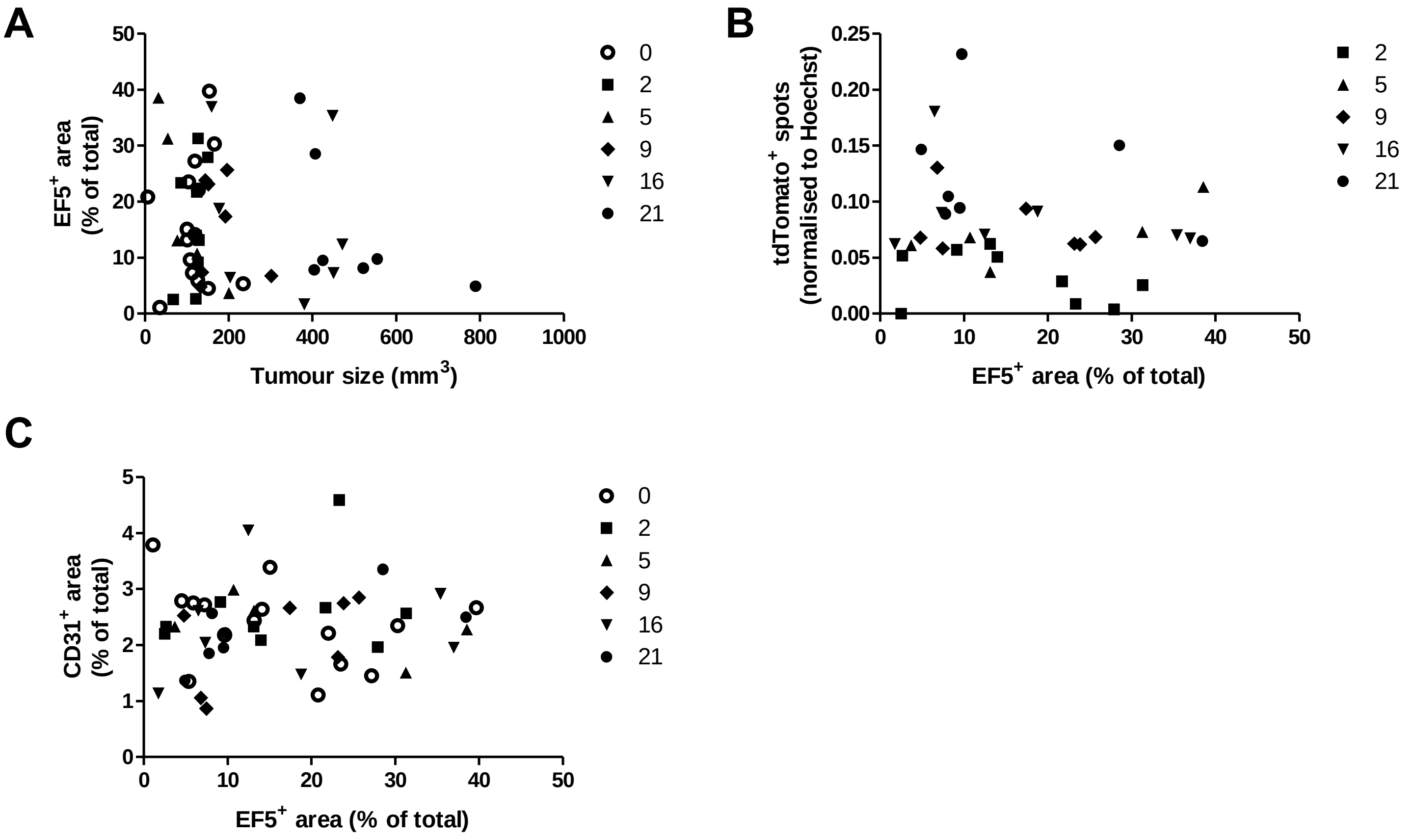


**Figure S3. Quantification of immunofluorescent staining of H1299-MR xenografts.** (A) Correlation between EF5 and tumours size (not significant). (B) Correlation between EF5 and tdTomato (not significant). (C) Correlation between EF5 and CD31 (not significant).

### Figure S4.

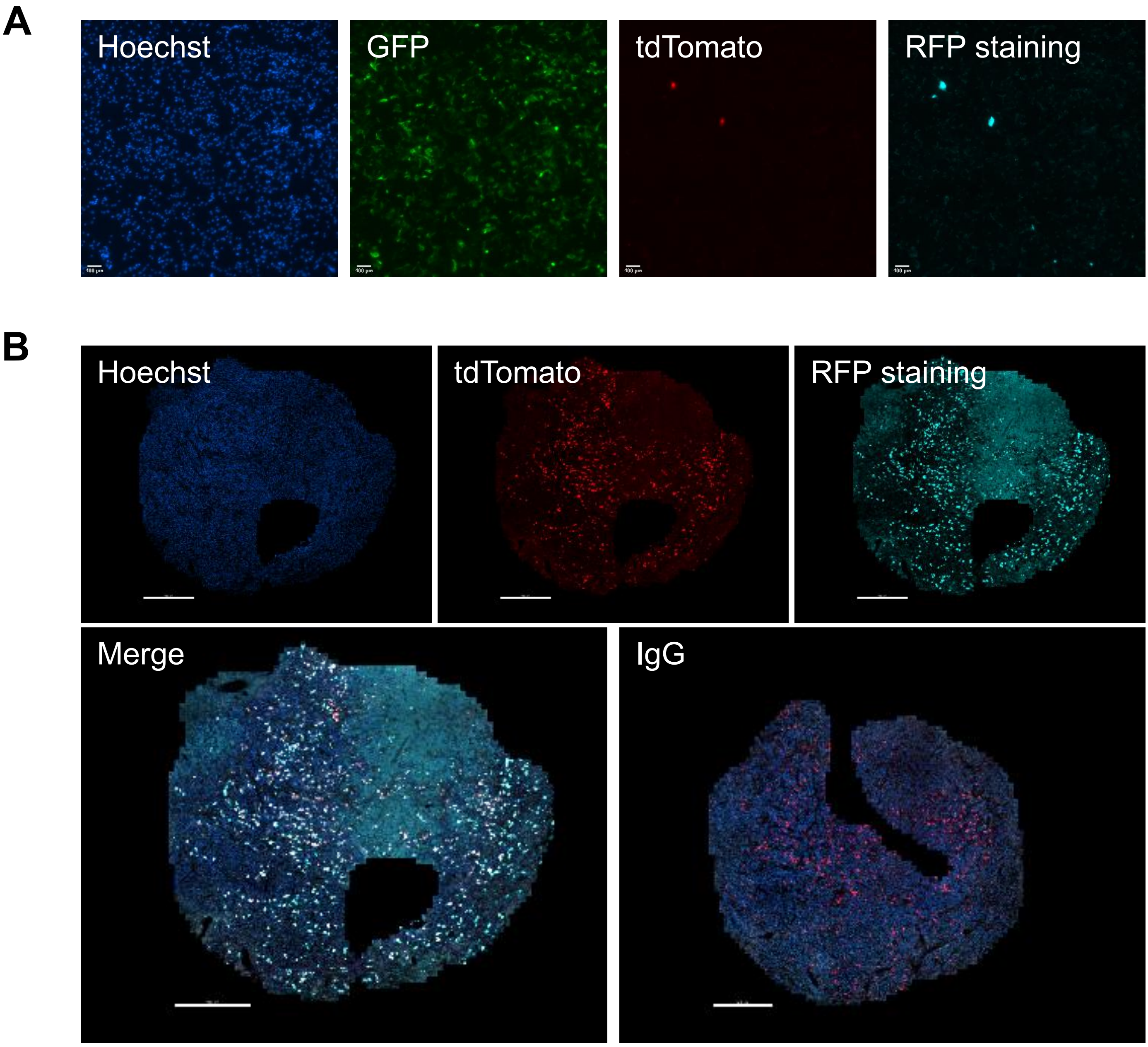


**Figure S4. Immunofluorescent RFP staining.** (A) Staining of GFP-positive cells, cultured *in vitro* with 100 μM DFO, confirming the antibody against RFP does not recognise GFP. (B) Micrographs of RFP staining and IgG control on xenograft sections. Scale bars A: 100 μm, B: 500 μm.

**Figure S5.**

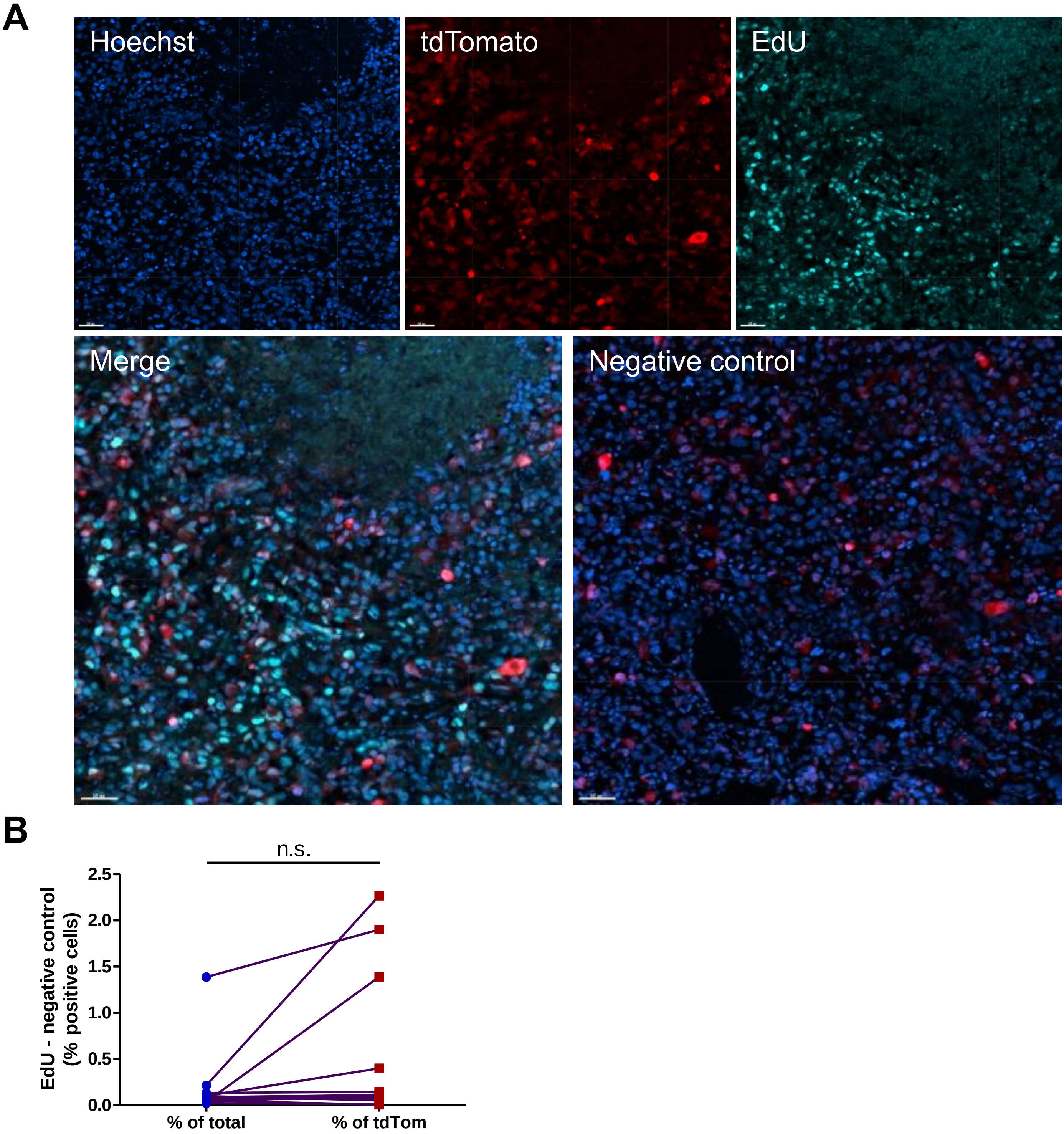


**Figure S5. EdU staining.** (A) Micrographs of xenograft sections stained for EdU and a negative control with only the AF-dye. Scale bars 50 µm. (B) Background staining as percentage positive cells of total (Hoechst) or tdTomato (n=13).

Figure S6.

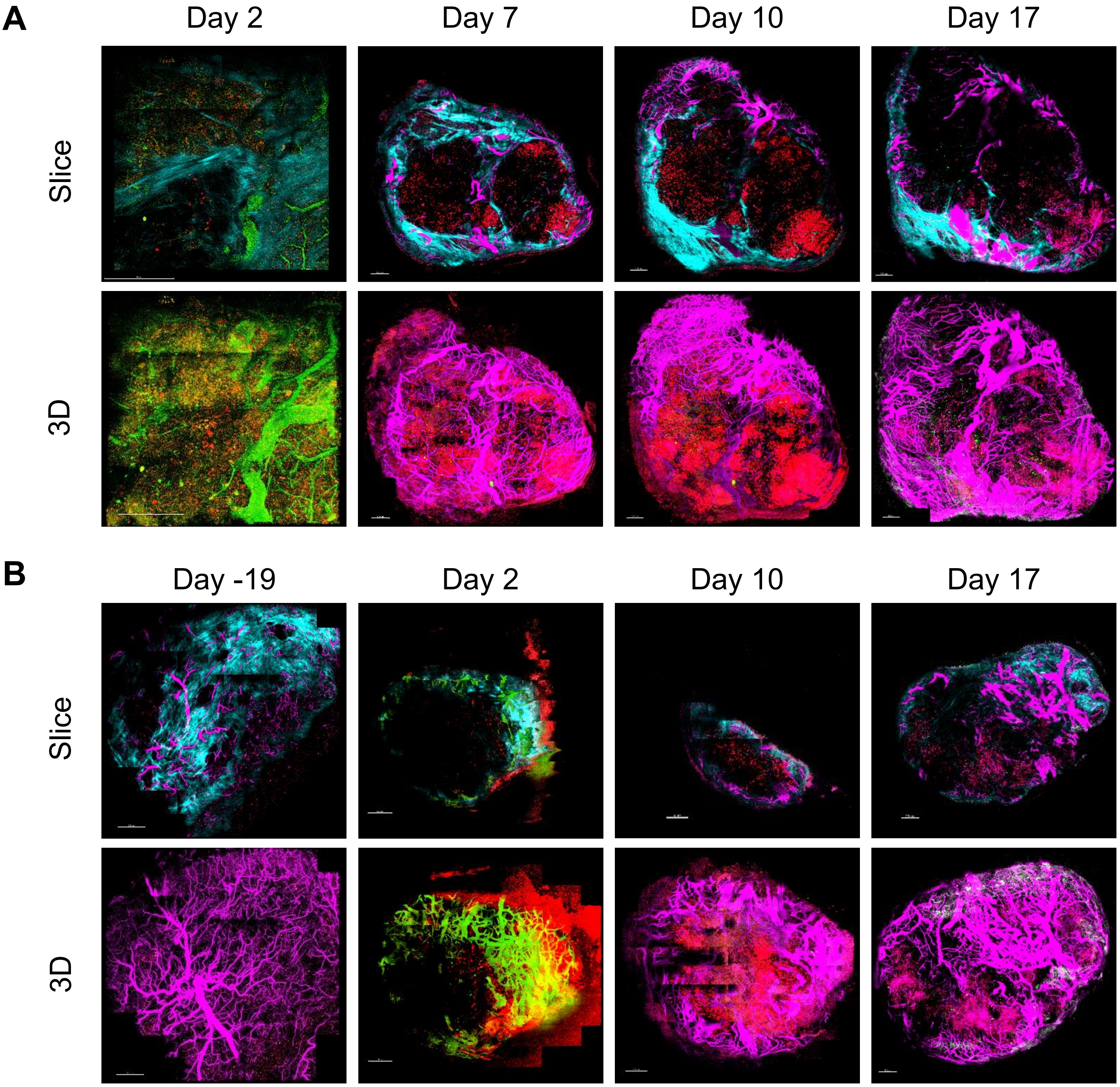
A
Day 2
Day 7
Day 10
Day 17
Slice
3D
B
Day -19
Day 2
Day 10
Day 17
Slice
3D

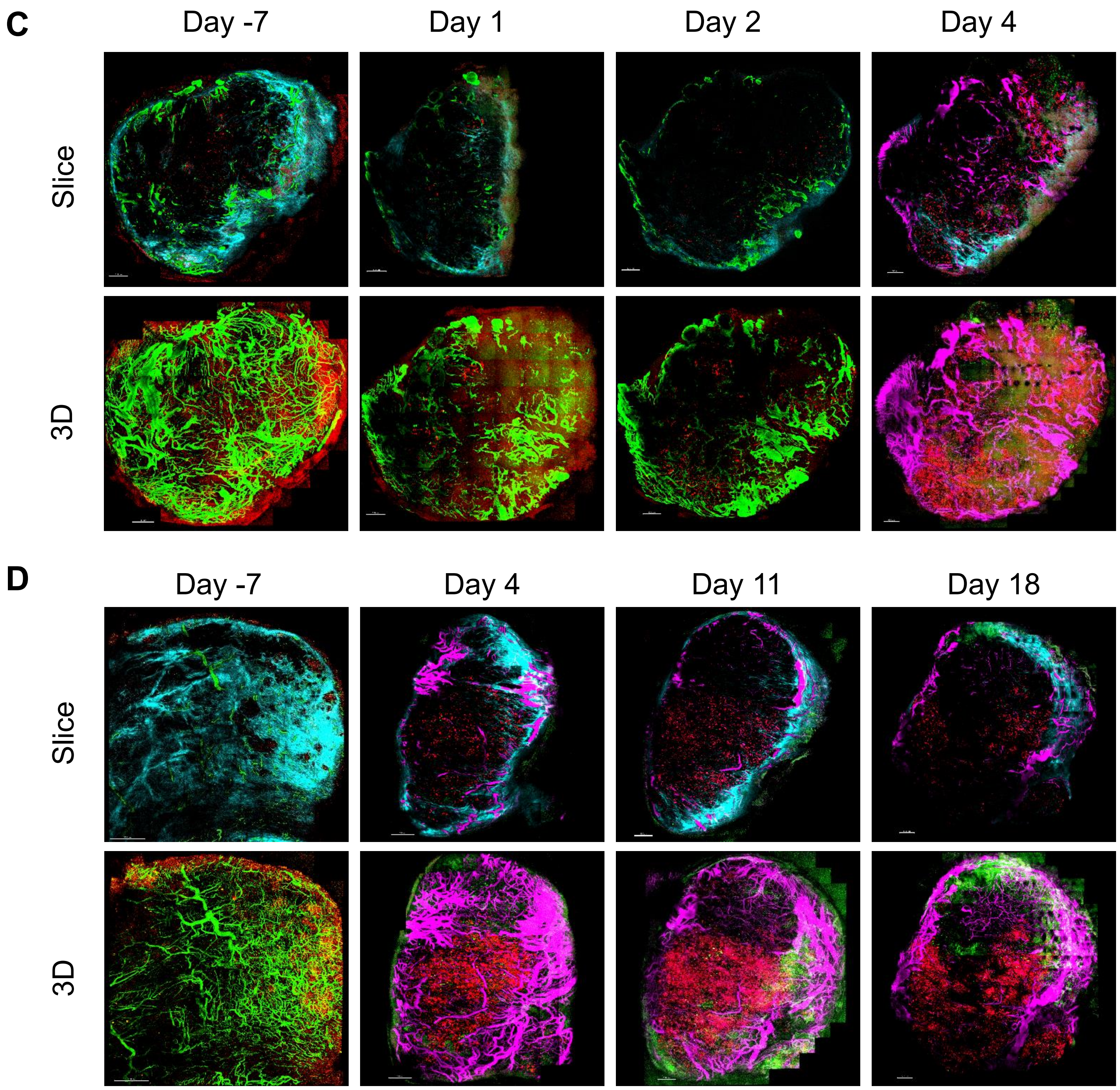


**Figure S6. H1299-MR xenografts in 4 mice were followed by intravital microscopy.** (A-D) Perfused vessels are shown in green (days -7, 1, and 2) or purple, tdTomato-positive cells in red, and collagen in cyan (second harmonic generation microscopy, top panels only). Days indicate the time after tamoxifen administration and scale bars are 500 μm. Channel arithmetics was applied using MATLAB to subtract GFP bleed through into the tdTomato channel and tdTomato bleed through into the Qtracker 705 channel (A-D).
